# Notch Signaling Reprograms Glial Lipid Metabolism to Promote Hypoxia Resistance

**DOI:** 10.64898/2026.06.26.734874

**Authors:** Yajuan Li, Qin Shuo, Anna Wang, Charlene Miciano, Allen Wang, Dan Zhou, Gabriel G. Haddad, Lingyan Shi

## Abstract

Hypoxia poses a major threat to the developing nervous system, where high metabolic demand is required to support brain growth, glial and neuronal maturation, and function. Although glial cells are essential for maintaining neural homeostasis under stress, how specific glial subtypes remodel metabolism to promote hypoxia tolerance remains poorly understood. Here, we identify a Notch-dependent lipid metabolic program in excitatory amino acid transporter 1 (Eaat1)-positive glia that supports hypoxia adaptation in the developing *Drosophila* larval brain. Using stimulated Raman scattering (SRS) microscopy combined with deuterium-labeled metabolic probes, we visualized substrate-specific metabolic activity in vivo at subcellular resolution. In control, non-adapted flies, we found that acute hypoxia markedly increased de novo lipogenesis in Eaat1-positive glia. In flies adapted to chronic hypoxia, Eaat1-positive glia exhibited a pre-programmed metabolic shift, characterized by reduced glucose-derived lipogenesis and enhanced acetate-derived lipid synthesis. Constitutive activation of Notch signaling in Eaat1-positive glia was sufficient to phenocopy this acetate-favored lipogenic state, suggesting that Notch promotes metabolic plasticity under oxygen-limited conditions. To define the transcriptional programs associated with this response, we performed single-nucleus RNA sequencing (snRNA-seq) of the developing *Drosophila* central nervous system and mapped Eaat-1expressing cell populations across hypoxia and Notch activation. Notch activation reshaped hypoxia-associated transcriptional responses and counteracted metabolic suppression caused by low oxygen. Together, our findings identify Eaat1-positiveglia as a metabolically adaptive glial population and reveal a conserved Notch-regulated mechanism that rewires lipid metabolism to support hypoxia tolerance in the developing brain. These results provide insight into glial metabolic strategies that may be relevant to hypoxia-associated neurological conditions, including neonatal hypoxic-ischemic brain injury and ischemic stroke.

## Introduction

Hypoxia is a fundamental physiological stress that challenges cellular metabolism, tissue homeostasis, and organismal survival ^1,2^. The nervous system is particularly vulnerable to oxygen deprivation because brain development requires high metabolic activity to support neurogenesis, glial maturation, synapse formation, and circuit assembly ^3,4^. Hypoxic stress during critical developmental windows can impair brain growth and cause long-term neurological dysfunction ^3,4^. In humans, oxygen deprivation contributes to conditions such as neonatal hypoxic-ischemic brain injury and ischemic stroke, where disrupted oxygen supply leads to profound metabolic stress, neuronal injury, and glial activation ^3–6^. Therefore, understanding how neural cells adapt to low oxygen is essential for defining the cellular mechanisms that protect the developing brain under hypoxic stress.

Cellular adaptation to hypoxia involves extensive metabolic reprogramming ^1,2^. The best-characterized response is the shift from oxidative phosphorylation toward anaerobic glycolysis, largely controlled by hypoxia-inducible factors (HIFs), which act as master regulators of oxygen homeostasis ^1,2,7^. Beyond glycolysis, emerging studies suggest that lipid metabolism is also an important component of the hypoxic response ^8,9^. Lipid metabolic remodeling may influence membrane homeostasis, energy storage, redox balance, oxidative stress buffering, and cell survival ^8,9^. However, how lipid metabolism is regulated in specific neural cell types during hypoxia, particularly in the developing brain, remains poorly understood. HIFs are well-established master regulators of hypoxic adaptation, while recent work supports a growing role for glial and neural lipid metabolism in stress responses.

Glial cells are central regulators of neural homeostasis. In addition to supporting neuronal metabolism and neurotransmitter balance, glia contribute to lipid handling, redox control, and tissue protection ^10–12^. These functions are especially important during hypoxia, when neurons face severe energetic and oxidative challenges ^13^. Despite their importance, the metabolic strategies used by defined glial subtypes to support hypoxia tolerance in vivo remain incompletely characterized.

The fruit fly, *Drosophila melanogaster*, has been a powerful genetic model system for dissecting complex biological systems in general and responses to hypoxia ^14^, in particular, owing to the high degree of conservation of disease-related genes and signaling pathways with humans ^15^. Previous genetic screens and long-term laboratory evolution experiments in *Drosophila* have identified several key signaling pathways, including Wnt and Notch, as crucial regulators of the hypoxic response ^16–18^. The Notch pathway is an evolutionarily conserved cell-cell communication system that plays a pivotal role in development, tissue homeostasis, and disease ^19,20^. Its activity is highly context-dependent, capable of promoting either oncogenic or tumor-suppressive functions ^21^. Beyond its roles in cell fate and proliferation, Notch signaling has also emerged as a key regulator of metabolic reprogramming in various biological systems ^22,23^. For instance, in mammalian cells, Notch has been shown to influence glucose metabolism by regulating the expression of glycolytic enzymes and mitochondrial respiration ^24^, impacting processes such as T-cell activation ^24^ and cancer cell proliferation ^25–27^.

Our previous study on *Drosophila* revealed that a particular subtype of cortex glia, distinguished by the presence of the excitatory amino acid transporter 1 (Eaat1), plays a vital role in ensuring survival under extreme hypoxic conditions ^17^. We demonstrated that the increased expression of Notch in Eaat1 cells is vital for the animal’s ability to withstand low oxygen conditions ^17,28^,. However, the precise molecular and cellular processes through which Notch signaling influences hypoxia resistance in Eaat1-positive cortical glia are still largely unidentified.

A major challenge in studying metabolic reprogramming *in vivo* has been the lack of tools to visualize metabolic reprogramming with high spatiotemporal resolution. Traditional methods like mass spectrometry-based metabolomics provide comprehensive data but destroy spatial context, while imaging techniques such as Positron Emission Tomography (PET) and Magnetic Resonance Spectroscopy (MRS) suffer from low spatial resolution, precluding single-cell analysis. The advent of stimulated Raman scattering (SRS) microscopy, a nonlinear optical imaging technique, has revolutionized the field ^27^. By coupling SRS with the use of small, non-perturbative vibrational tags, such as deuterium (D), it is now possible to trace the fate of labeled metabolites such as deuterated glucose, acetate, amino acids or heavy water into newly synthesized macromolecules *in situ*, providing a dynamic, volumetric map of metabolic activity at the subcellular level ^30–32^.

In this study, we leveraged the power of *Drosophila* genetics, flies adapted to chronic hypoxia, and SRS microscopy as well as single-nucleus RNA sequencing (snRNA-seq) to investigate the mechanisms of hypoxia adaptation in the brain. By SRS imaging, we mapped the spatiotemporal dynamics of lipogenesis in response to hypoxia. By utilizing snRNA-seq, we mapped the cellular composition of the developing *Drosophila* CNS and investigated how different neural cell types transcriptionally respond to hypoxia. We specifically focus on the Eaat1-positive glial cells to understand the molecular basis of Notch-mediated hypoxia tolerance. This study provides a detailed analysis of the cellular and molecular mechanisms involved in the hypoxic response of a developing brain, highlighting the distinctive metabolic strategies glial cells employ to manage oxygen deprivation and ensure neuroprotection.

## Results

### Hypoxia Induces De Novo Lipogenesis in Eaat1 Glial Cells

To investigate the metabolic response to hypoxia in *Drosophila* larval brain, we first sought to identify the primary sites of lipid metabolism. We integrated two photon fluorescence (TPF) with SRS microscopy to image endogenous lipids (targeting the CH□ stretching vibration at 2850 cm^-1^) in the larval brain, where *elav*-, *repo*-, *Eaat1*-GAL4 was used to drive the expression of a nuclear GFP reporter in pan-neuron, glia and Eaat1 positive cells, respectively (Fig. 1A, B). Our imaging revealed highly abundant lipid droplets in the cytoplasm of glial cells, particularly Eaat1-positive glial cells (Eaat1-glia), which are glial subtype ensheathed the neuronal cell bodies in the brain cortex, whereas neurons contained little lipid (Fig. 1B). This indicates that Eaat1-glia are a major hub for lipid storage in the brain.

**Figure 1.**
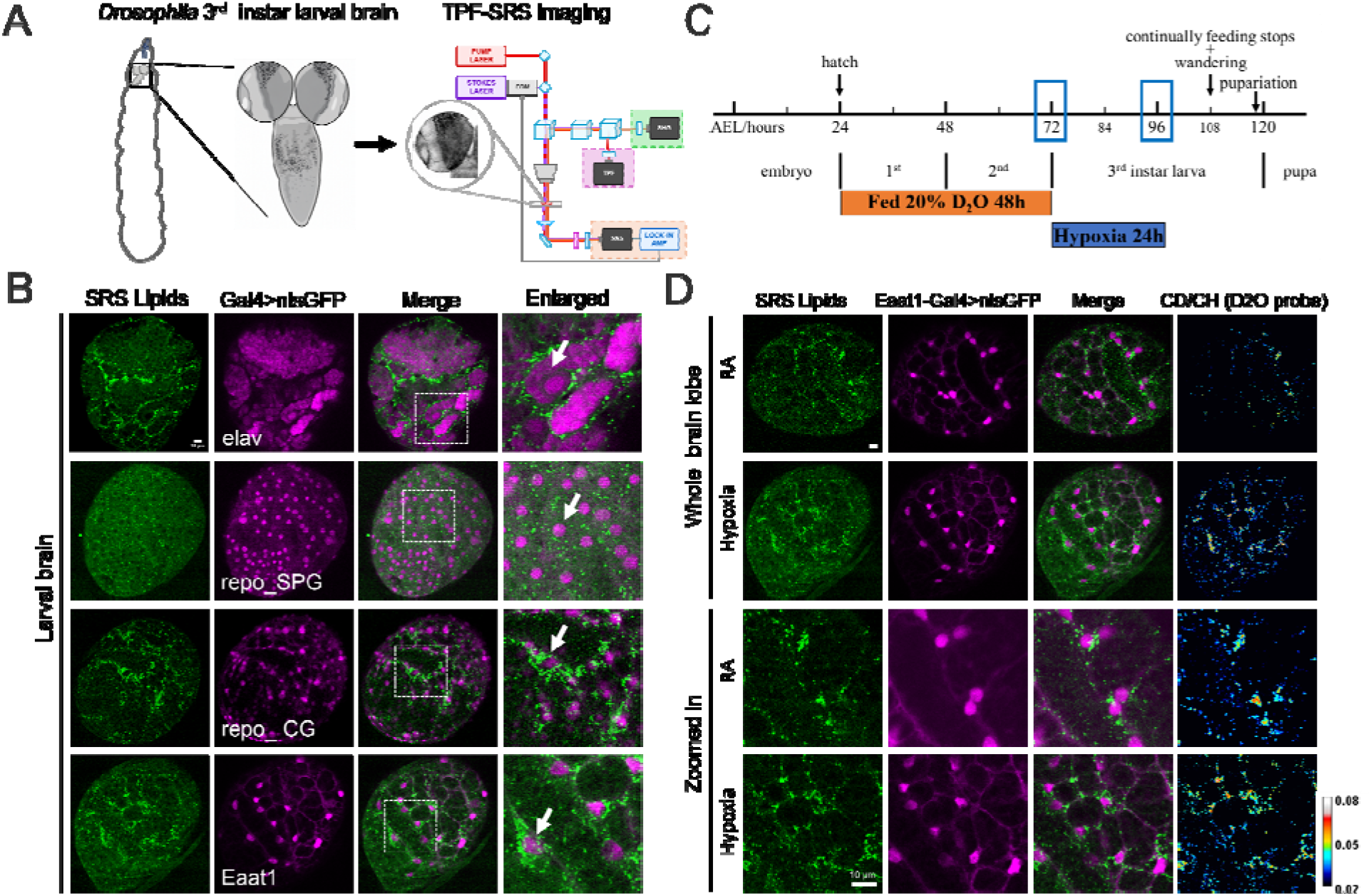
Hypoxia increases de novo lipid synthesis in Eaat1-positive glia. **(A)** Schematic of two photon fluorescence (TPF)-SRS imaging of *Drosophila* third-instar larval brain, consisting of two lobes and a ventral nerv cord. **(B)** Representative SRS image of endogenous lipids (green, 2850 cm□^1^) overlaid with a TFP image of Eaat1-glia marked by nuclear GFP (nlsGFP, magenta). Lipids are predominantly localized within the glial cytoplasm, particularly in Eaat1-glia. In contrast, lipids are barely found in elav-nlsGFP expressing neurons. SPG (subperineurial glia), CG (cortex glia). Scale bar, 10 μm. The subcellular distribution of lipids was indicated by the white arrows in the enlarged images. **(C)** Experimental scheme for metabolic labeling and hypoxia treatment. First-instar larvae were fed D□O-containing food for 48 hours and then treated by either room air (RA, 21% O□) or hypoxia (3.5% O□) for 24 hours. **(D)** Representative CD/CH ratiometric SRS images of newly synthesized lipids in larval brains from room air (RA) and hypoxia. The CD/CH ratio was calculated by the intensity at 2140 cm□^1^/2850 cm□^1^. The results show hypoxia leads to a marked increase in lipid synthesis in Eaat1-glia. Scale bar, 10 μm.

To measure the rate of new lipid synthesis, we fed larvae a diet containing heavy water (D□O), a general metabolic tracer. The deuterium (D) from D□O is incorporated into macromolecules during their synthesis, and the resulting C-D bonds can be specifically imaged by SRS microscopy in the cell-silent spectral region (∼2100-2200 cm^-1^). By calculating the CD/CH ratio at 2140 cm□^1^/2850 cm□^1^, ratiometric images can be generated to map and quantify newly synthesized lipids at subcellular resolution. After feeding larvae D□O for 48 hours and then exposing them to either room air (RA, 21% O□) or hypoxia (3.5% O□) for the following 24 hours (Fig. 1C), we observed a dramatic increase in CD/CH ratio within Eaat1-glia of hypoxia-treated brains compared to RA controls (Fig. 1D). This result demonstrates that hypoxia trigger a significant upregulation of de novo lipogenesis specifically within Eaat1-glia subtype.

### Hypoxia-Adapted Flies Display a Reprogrammed Lipid Metabolic State

We next investigated how lipid metabolism is altered in flies that are adapted to chronic hypoxia. This is a unique *Drosophila* line, termed low-oxygen adapted flies (AF), which has been selected for survival and reproduction in an increasingly hypoxic environment for over 300 generations ^16^. We compared the rate of lipid synthesis in AF with that of the control naïve flies (NF) along different developmental stages under RA condition. SRS imaging of D□O-labeled lipids revealed that AF exhibited a significantly lower basal rate of lipogenesis throughout development compared with NF (Fig. 2A, B). However, when subjected to the hypoxic challenge, AF showed a remarkable increase in lipid synthesis compared with NF (Fig. 2C). These findings suggest that chronic adaptation to hypoxia involves a fundamental reprogramming of lipid metabolism, resulting in a lower basal lipid metabolic rate and an enhanced response to acute oxygen deprivation. This pre-adapted state may be more efficient and sustainable in a constantly low-oxygen environment.

**Figure 2.**
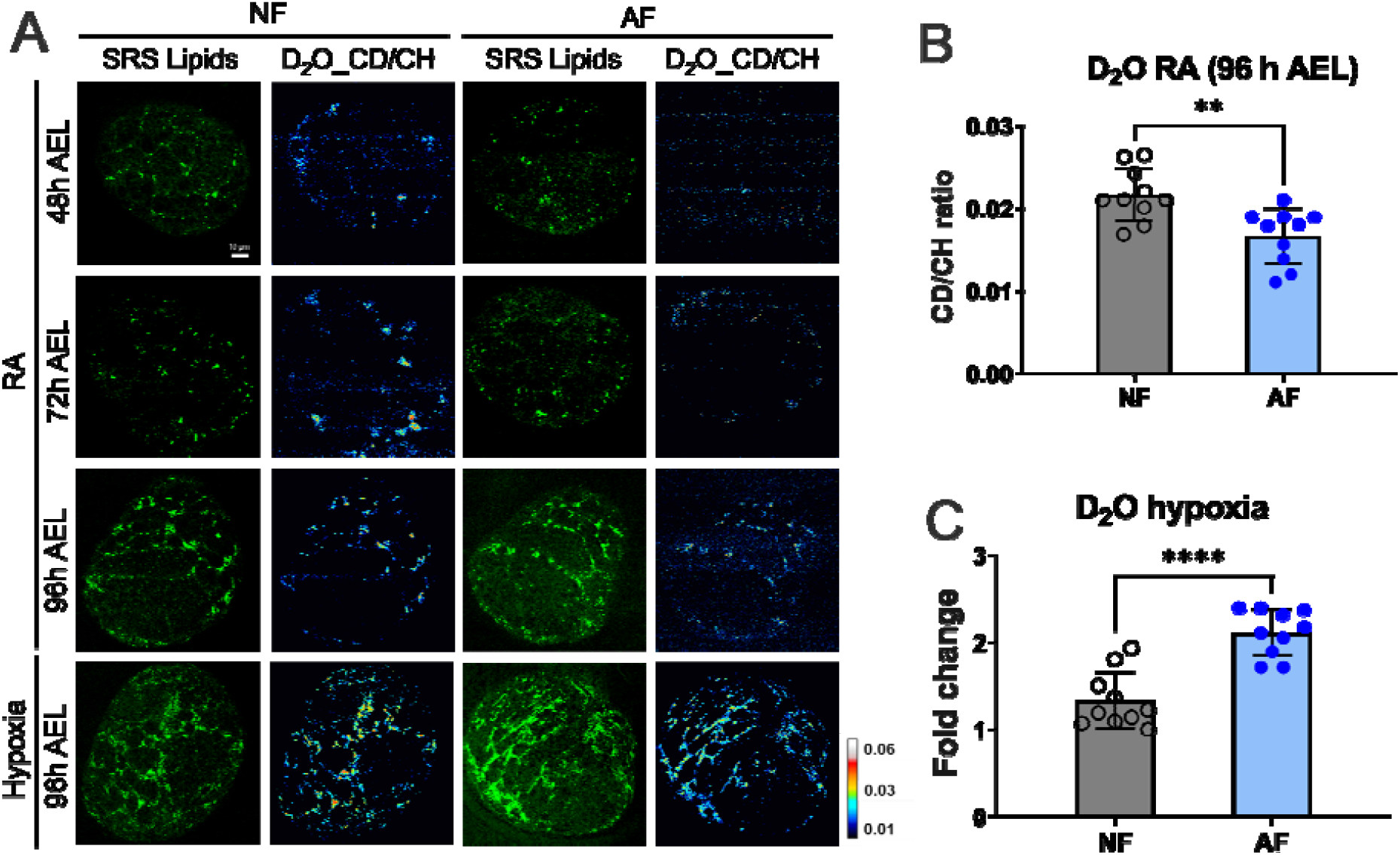
Hypoxia-adapted flies exhibit reduced basal and hypoxia-induced lipogenesis. **(A)** Representative CD/CH ratiometric SRS images of newly synthesized lipids in larval brains of naïve flies (NF) and low-oxygen adapted flies (AF) under room air (RA), showing the increased lipid synthesis during development and hypoxia. **(B)** Quantification of the basal lipid synthesis rate by CD/CH ratio, showing a significant reduction in AF compared to NF at 96 h after egg laying (AEL). **(C)** Quantification of hypoxia-induced lipogenesis based on the CD/CH ratio, shown as fold change in hypoxia-treated samples relative to room air (RA) controls. AF show a significant increase in lipid synthesis upon hypoxic exposure compared with NF. Data was shown as mean ±SD, **p < 0.01, ****p < 0.0001; Student’s *t*-test; n=10 flies.

### A Metabolic Switch from Glucose to Acetate Underlies Hypoxia Adaptation

Lipids can be synthesized from various carbon sources, with glucose and acetate being two major precursors. To dissect the specific metabolic pathways altered in AF, we used SRS microscopy to trace the incorporation of deuterated glucose ([D□]-glucose) and deuterated acetate ([D□]-acetate) into lipids. Under RA condition, we found that NF primarily utilized glucose for lipid synthesis, whereas AF showed a marked preference for acetate (Fig. 3A-3C). This indicates a shift in the primary carbon source for lipogenesis as a result of adaptation to chronic hypoxia. Upon hypoxic exposure, this metabolic divergence was further amplified. AF exhibited reduced glucose-derived lipogenesis, likely due to impaired aerobic glucose metabolism under low-oxygen conditions. By contrast, acetate-derived lipogenesis was significantly increased in AF under hypoxia. Fold-change analysis further confirmed this substrate-specific rewiring: the hypoxia-to-room-air ratio decreased for glucose-derived lipid labeling but increased for acetate-derived lipid labeling. These results indicate that hypoxia shifts AF lipid synthesis away from glucose-dependent carbon input and toward acetate utilization, suggesting that acetate becomes a preferred substrate for maintaining lipogenesis under oxygen-limited conditions (Fig. 3D, 3E). This demonstrates a dynamic and adaptive metabolic switch: hypoxia-adapted flies not only rely on acetate at baseline but also enhance this pathway to fuel lipogenesis during periods of low oxygen stress.

**Figure 3.**
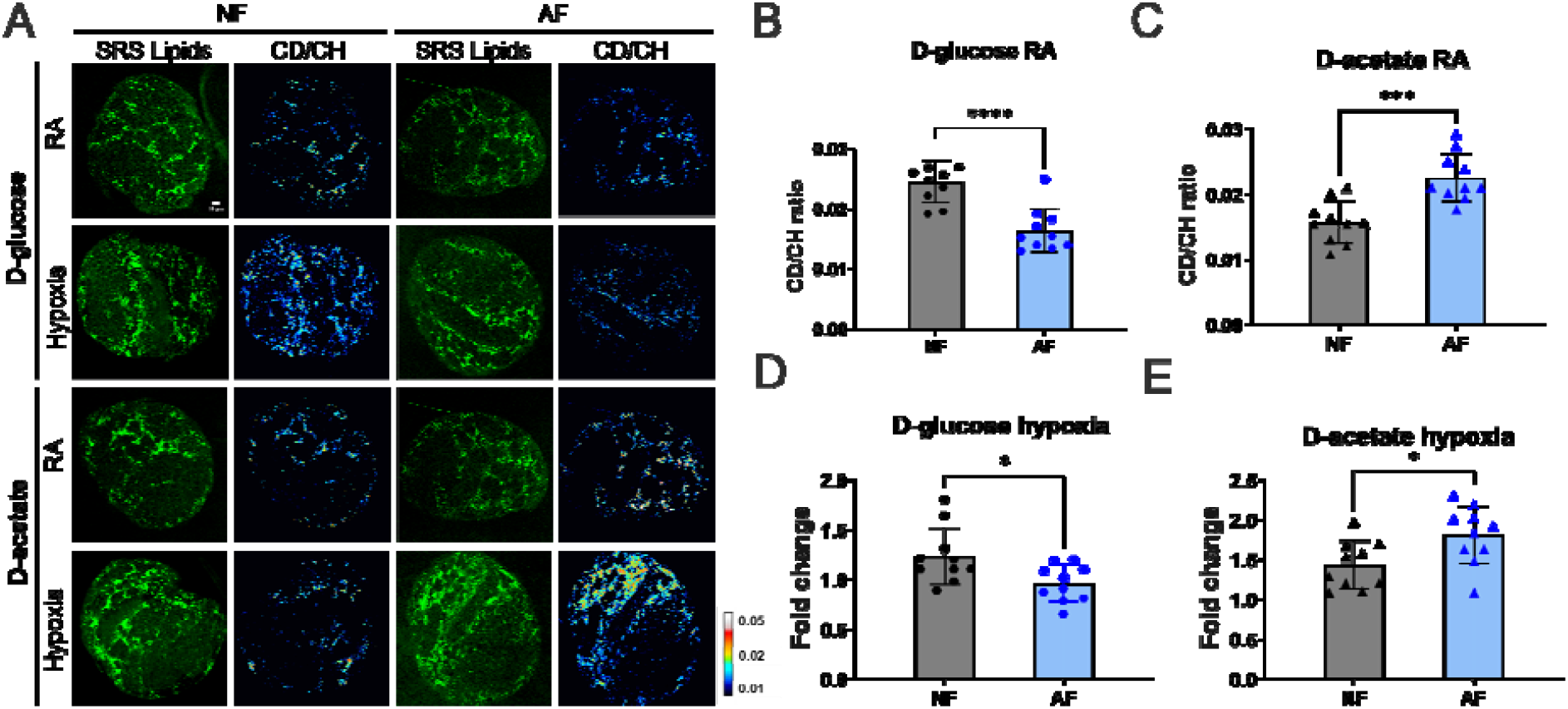
Hypoxia-adapted flies shift lipid synthesis from glucose to acetate utilization. **(A)** Representative SRS images of newly synthesized lipids derived from either [D□]-glucose or [D□]-acetate in NF and LOF brains under room air (RA) and hypoxia. **(B)** Quantification showing that RA condition the basal lipid synthesis from glucose is reduced in AF compared to NF. **(C)** Quantification showing that basal lipid synthesis from acetate is increased in AF compared to NF. **(D–E)** Fold-change quantification of substrate-derived lipogenesis in AF under hypoxia relative to room air. Glucose-derived lipogenesis was decreased under hypoxia (D), whereas acetate-derived lipogenesis was significantly increased (E), indicating a hypoxia-induced shift from glucose-to acetate-dependent lipid synthesis in AF. Data was shown as mean ±SD, *p < 0.05, ***p < 0.001; Student’s *t*-test; n=10 flies.

### Notch Signaling Activation Mimics the Metabolic and Survival Phenotype of Hypoxia-Adapted Flies

Previous genetic screens implicated the Notch signaling pathway in Eaat1-glial cells mediated hypoxia tolerance ^13,16^. To test whether Notch signaling could regulate the observed metabolic switch, we constitutively activated the pathway specifically in Eaat1-glia by overexpressing the Notch Intracellular Domain (*Eaat1*>*NICD*). Remarkably, under room air condition, *Eaat1*>*NICD* flies phenocopied the metabolic signature of AF. They exhibited a significant reduction in glucose-derived lipogenesis and a corresponding increase in acetate-derived lipogenesis compared to control flies *(Eaat1*>*lacZ*) (Fig. 4A-4C). Notably, when exposed to a hypoxic environment (3.5% O□), *Eaat1*>*NICD* flies exhibited a more pronounced and significant shift in metabolism towards acetate utilization compared to *Eaat1*>*lacZ* controls (Fig. 4D, 4E). These results strongly suggest that Notch activation is sufficient to orchestrate the metabolic shift from glucose to acetate utilization in Eaat1-glia.

**Figure 4.**
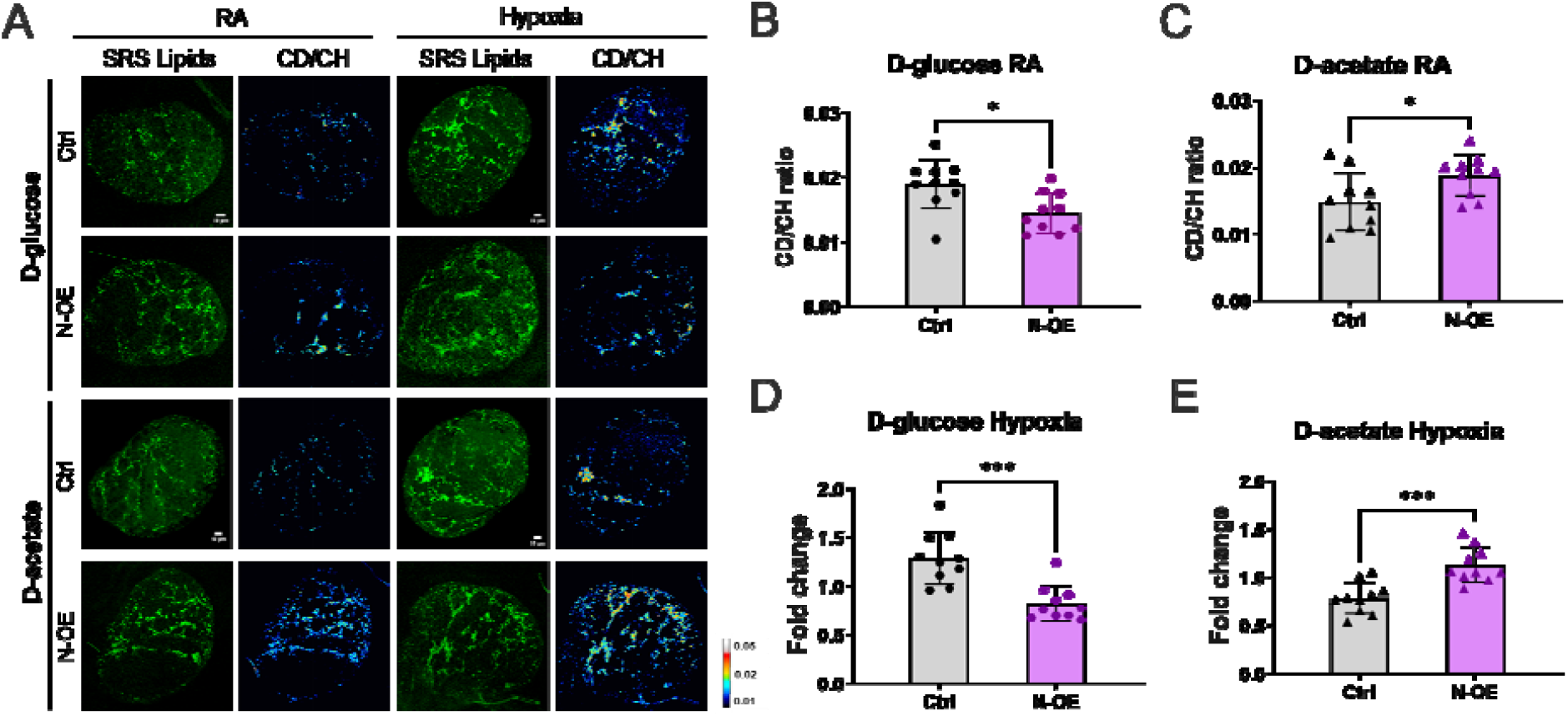
Notch activation phenocopies the metabolic switch of hypoxia-adapted flies. **(A)** Representative SRS images of newly synthesized lipids from [D□]-glucose or [D□]-acetate in *Eaat1*>*UAS*-*lacZ* (Ctrl) and *Eaat1*>*NICD* (N-OE) flies under room air (RA). **(B–E)** Quantification of glucose- and acetate-derived lipogenesis upon Notch activation. (B, C) CD/CH ratios show that Notch activation decreases glucose-derived lipogenesis (B) but increases acetate-derived lipogenesis (C) under room air. (D, E) Fold-change analysis under hypoxia relative to room air shows reduced glucose-to-lipid flux (D) and enhanced acetate-to-lipid flux (E) following Notch activation. These results suggest that Notch activation promotes an AF-like metabolic shift from glucose- to acetate-dependent lipid synthesis. Data was shown as mean ±SD, *p < 0.05, ***p < 0.001; Student’s *t*-test; n=10 flies.

### Single-Nucleus Profiling Identifies Eaat1-Positive Glial Populations under Hypoxia and Notch Activation

To identify the Eaat1-expressing cell populations that may mediate Notch-dependent hypoxia adaptation, we performed snRNA-seq on dissected 3^rd^ instar larval brains in four experimental conditions: room air control, room air with Notch overexpression (N-OE), hypoxia control, and hypoxia with N-OE. After stringent quality control, we obtained a total of 54,224 high-quality nuclear transcriptomes across all experimental conditions. Unsupervised clustering and UMAP visualization revealed 28 distinct clusters, which were assigned to major cell classes based on the expression of canonical marker genes (Figure 5A, Table 1). We identified neural stem cells (neuroblasts, NBs) marked by *mira* and *dpn*, and their progeny, ganglion mother cells (GMCs), expressing *erm* and *hey*. A large and diverse population of neurons was identified by markers such as *nSyb* and *elav*. We also resolved multiple glial subtypes, including astrocyte-like glia (*Eaat1, gs2*), ensheathing glia (*moody*), cortex glia (*wrapper*), and perineurial glia (*moody*). Other identified cell types included hemocytes (*Hml*) and cells of the tracheal system (*trh*) (Figure 5B). Together, these data define the cellular context for analyzing hypoxia- and Notch-dependent responses in Eaat1-expressing cells.

**Table 1.**
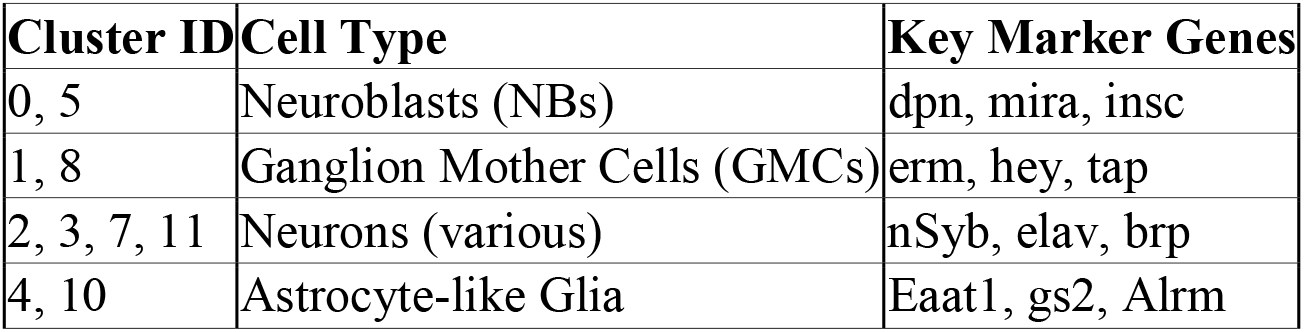
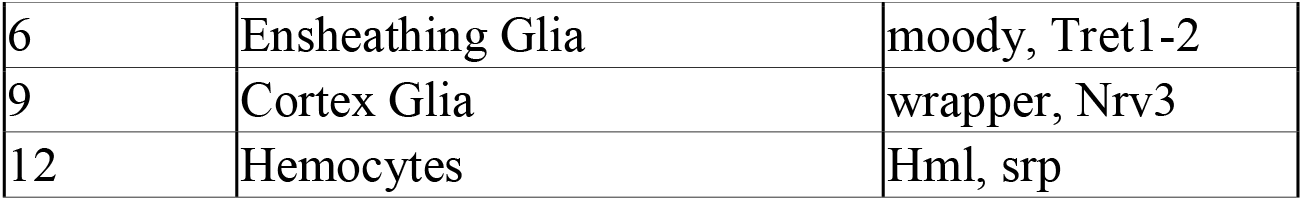
Major Cell Clusters and Key Marker Genes.

**Figure 5.**
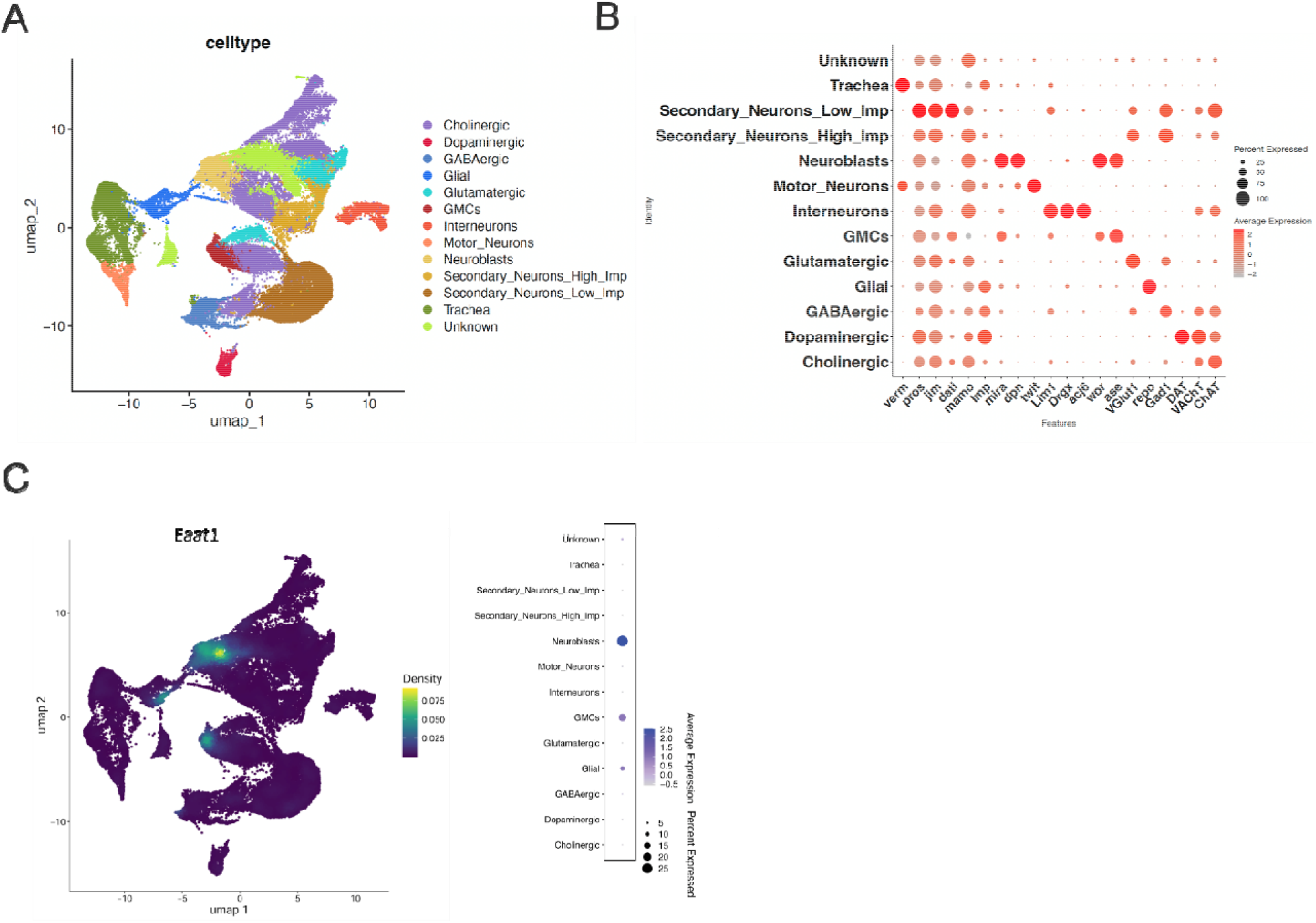
Single-nucleus transcriptomic profiling reveals glial-specific responses and metabolic reprogramming under hypoxia. **(A)** Uniform Manifold Approximation and Projection (UMAP) of single-nucleus RNA sequencing data from developing *Drosophila* brains, colored by annotated cell types. Major neuronal and non-neuronal populations are identified, including cholinergic, dopaminergic, GABAergic, glutamatergic neurons, glia, mushroom body neurons, interneurons, and other minor cell populations. **(B)** Dot plot showing expression of canonical marker genes across annotated cell types. Dot size represents the percentage of cells expressing each gene, and color intensity indicates average expression level. These markers validate cell-type identities, including neuronal subtypes and glial populations. **(C)** Feature plot showing the distribution and expression density of the Eaat1 marker across the UMAP embedding. High Eaat1 expression is restricted to neuroblast, ganglion mother cell (GMC) and glia.

Previous work has implicated Notch signaling in Eaat1-positive cells as a critical mediator of hypoxia tolerance in *Drosophila* ^13,33^. We identified 1,159 Eaat1-expressing nuclei, which were predominantly located in three distinct clusters: 7, 14, and 15. These clusters likely represent distinct Eaat1-expressing cell populations of neuroblast, ganglion mother cell (GMC) and glia (Figure 5C). We next examined the relative contribution of these glial subpopulations across our four experimental conditions: room air control, room air with Notch overexpression (N-OE), hypoxia control, and hypoxia with N-OE. Analysis of cell proportions revealed dynamic changes in the composition of the different cell types in response to both hypoxia and Notch activation (Fig. S1A), including the changes of relative abundance of cells within clusters 7, 14, and 15 (i.e., Eaat1-positive cells). These compositional shifts suggest that environmental and genetic perturbations can alter the cellular landscape of the brain, potentially reflecting adaptive or pathological responses.

### Notch Activation Counteracts Hypoxia-Associated Suppression of Metabolic Expression Programs

To define the transcriptional programs associated with Notch-dependent responses, we performed Gene Set Enrichment Analysis (GSEA) on the combined transcriptomes of Eaat1-positive cells from clusters 7, 14, and 15. This analysis indicated that hypoxia alters pathways related to oxidative phosphorylation, fatty acid metabolism, cholesterol homeostasis, and inflammatory signaling, suggesting consistent with broad metabolic and stress-response remodeling in Eaat1-expressing cells (Figure S1B).

We next focused on Eaat1-glial cells to examine metabolic transcriptional responses more directly. First, we assessed the impact of hypoxia alone by comparing control flies in hypoxia versus room air. This comparison revealed a significant downregulation of genes associated with oxidative phosphorylation (Fig. 6A). This finding is consistent with the established cellular response to oxygen deprivation, where cells often suppress aerobic respiration to conserve resources and reduce the production of reactive oxygen species. Next, we investigated the effect of activating the Notch pathway under hypoxic stress by comparing the hypoxia N-OE group to the hypoxia control group. Remarkably, GSEA revealed that Notch overexpression led to a significant upregulation of the oxidative phosphorylation pathway, effectively counteracting the suppressive effect of hypoxia (Fig. 6A). These findings imply that activating Notch might grant hypoxia tolerance by improving or restoring mitochondrial respiratory function in Eaat1-positive glial cells. We utilized two-photon fluorescence imaging of NADH and Flavin to assess the redox ratio (NADH/Flavin), an indicator of cellular redox state and metabolic activity, under various conditions. The results reveal that the reduction in the redox ratio caused by hypoxia was reversed by overexpressing Notch in Eaat1-positive cells (Fig. 6B). This data corroborates the snRNA-seq findings.

**Figure 6.**
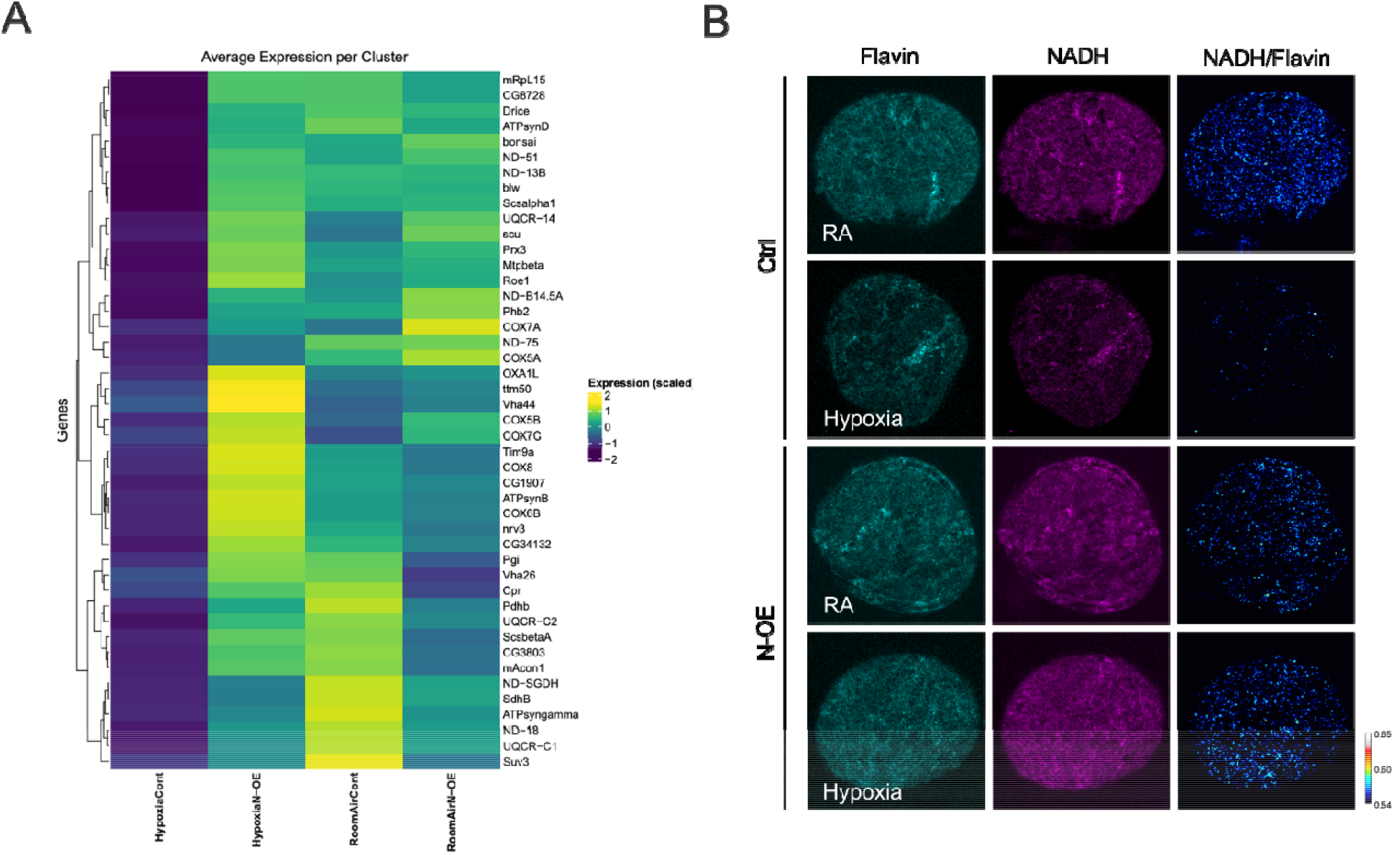
Hypoxia and Notch signaling modulate mitochondrial metabolism and redox state in *Drosophila* brain. **(A)** Heatmap of scaled gene expression showing average expression levels of mitochondrial and metabolic genes across experimental conditions (hypoxia versus room air; control versus N-OE). Genes associated with oxidative phosphorylation, electron transport chain complexes, and mitochondrial function exhibit coordinated changes, indicating metabolic reprogramming under hypoxia and genetic perturbation. **(B)** Representative fluorescence images of NADH and flavin in control and N-OE larval brains under RA and hypoxia conditions. Notc overexpression revers the reduction in redox balance under hypoxia treatment.

## Discussion

This study identifies a glial metabolic response to hypoxia in the developing *Drosophila* brain, characterized by lipid accumulation, altered substrate utilization for lipid synthesis, and change in cellular metabolic state. By applying substrate-resolved stimulated Raman scattering (SRS) microscopy in vivo, we visualized lipid metabolic activity with cell-type and subcellular resolution. Our findings show that Eaat1-positive glia exhibit a robust lipogenic response under hypoxia and hypoxic adaptation, and that activation of Notch signaling in these glia is sufficient to induce lipid remodeling and metabolic features associated with the hypoxic state. Together, these results support a model in which Notch signaling contributes to glial metabolic plasticity during hypoxia.

A central finding of this work is that hypoxia does not simply increase lipid accumulation uniformly across the brain but instead induces a selective and substrate-dependent remodeling of lipid metabolism in Eaat1-positive glia. Using deuterium- and isotope-labeled metabolic probes, we observed changes in the contribution of different carbon sources to newly synthesized lipids. In particular, hypoxic adaptation was associated with a shift toward acetate-derived lipid synthesis, suggesting that glial cells can reorganize substrate utilization under reduced oxygen availability. This substrate flexibility may provide an adaptive metabolic strategy when canonical energy and biosynthetic pathways are constrained by hypoxia. However, because the current study does not directly test whether acetate-driven lipogenesis is required for hypoxia survival, we interpret this metabolic switch as a candidate adaptive mechanism rather than a proven causal determinant of hypoxia resistance.

Our data also link lipid metabolic remodeling with broader changes in mitochondrial and redox-associated states. The snRNA-seq analysis revealed transcriptional changes related to oxidative phosphorylation, mitochondrial function, and lipid metabolism, particularly within glial populations. These transcriptional features are consistent with the SRS imaging results and suggest that lipid synthesis, mitochondrial metabolism, and cellular redox state are coordinated during the hypoxic response. Rather than representing isolated metabolic outputs, these changes may reflect an integrated glial program that balances biosynthesis, energy metabolism, and stress adaptation. Future experiments that directly perturb acetate metabolism, fatty acid synthesis, mitochondrial activity, or redox regulation will be important to determine the causal order among these processes.

Our findings further implicate Notch signaling as an upstream regulator of this glial metabolic state. Notch is best known for its roles in cell fate specification and developmental patterning, but increasing evidence supports additional roles in regulating cellular physiology and metabolism. In this study, Notch activation in Eaat1-positive glia was sufficient to induce lipid accumulation and metabolic remodeling resembling features of the hypoxic response. These results suggest that Notch may link environmental stress signals to metabolic reprogramming in a defined glial population. At the same time, it remains important to distinguish metabolic regulation from potential changes in cell fate or cell composition. Therefore, the Notch gain-of- function phenotype should be interpreted together with glial identity markers and snRNA-seq cluster annotations to ensure that the observed metabolic changes arise from altered cellular state rather than a major shift in cell identity or population composition.

The cell-type specificity of this response is particularly important. Our imaging and genetic analyses indicate that Eaat1-positive glia are a major site of hypoxia-induced lipid remodeling. This observation supports the idea that different brain cell types adopt distinct metabolic strategies under oxygen limitation. Eaat1-positive glia may serve as metabolic responders that adjust lipid synthesis and storage in response to hypoxic stress. The functions of the accumulated lipids remain to be determined. They may contribute to membrane remodeling, lipid storage, or stress buffering, but these possibilities require direct functional testing. Similarly, although lipid droplets have been implicated in oxidative stress responses in other contexts, our current data do not directly establish that lipid accumulation in Eaat1-positive glia reduces oxidative damage or preserves energy reserves during hypoxia.

The relationship between glial lipid remodeling and hypoxia tolerance is therefore best viewed as a working model. Previous studies have shown that hypoxia tolerance can involve coordinated genetic and metabolic adaptation, supporting the idea that organismal resistance emerges from multi-layered physiological programs. Our study extends this concept by identifying a cell-type-specific metabolic phenotype within the brain and by providing imaging evidence for substrate-specific lipid synthesis during hypoxia. However, additional functional experiments will be needed to determine whether blocking acetate utilization, fatty acid synthesis, or Notch-induced lipid remodeling is sufficient to impair hypoxia adaptation or survival.

These findings also provide a framework for considering conserved principles of glial metabolic plasticity, but the broader relevance to mammalian systems should be interpreted cautiously. Glial lipid metabolism, mitochondrial remodeling, and hypoxia responses are important across species, yet the specific role of Eaat1-positive glia in Drosophila cannot be directly equated with mammalian glial cell types without additional comparative evidence. Thus, rather than making a direct clinical claim, our study highlights a potentially conserved biological principle: glial cells can undergo regulated metabolic remodeling in response to oxygen limitation, and developmental signaling pathways such as Notch may participate in coordinating this response.

In summary, our study combines substrate-resolved SRS imaging, transcriptomic profiling, and glial-specific genetic manipulation to identify Notch-associated lipid metabolic remodeling in Eaat1-positive glia during hypoxia. The strongest evidence supports a model in which Notch activation is sufficient to promote a hypoxia-like glial metabolic state involving lipid accumulation, altered substrate use, and changes in OXPHOS/redox-associated programs. While future studies are needed to establish direct functional causality between the metabolic switch and hypoxia resistance, our work reveals glial lipid metabolic plasticity as an important component of the brain’s response to hypoxic stress.

## Methods

### *Drosophila* Stocks and Genetics

*Drosophila melanogaster* were raised on standard cornmeal-yeast-agar medium at 25°C on a 12:12 hour light:dark cycle. The AF and NF strains were generated from the previously described populations obtained from experimental evolution under normoxic or hypoxic conditions ^16,17^. The wild-type (WT) control strain (NF) was generated from the control population that were maintained in normoxia/room air condition, and the low-oxygen-adapted flies (AF) were derived from the hypoxia-adapted populations that were evolving under chronic hypoxic conditions for over 300 generations. The GAL4/UAS system was used for cell-type-specific gene expression. The *Eaat1-GAL4* driver line was used to target expression to a specific subtype of cortex glia. The following UAS lines were obtained from the Bloomington Drosophila Stock Center: *Elav-GAL4* (pan neuronal expression), *UAS-lacZ* (overexpressing lacZ as genetic control), *UAS-nlsGFP* (nuclear localized Green Fluorescent Protein). The *UAS-NICD* (constitutively active Notch Intracellular Domain) stock was provided by Dr. J. Posakony. Genetic crosses were set up to generate flies of the desired genotypes.

### Hypoxia Treatment

For hypoxia experiments, third-instar larvae were placed in a computer-controlled hypoxia chamber (Coy Laboratory Products) and exposed to a gas mixture of 3.5% O□ and 96.5% N□ for 24 hours. Room air controls were maintained in parallel in ambient air (21% O□). Each experiment consists of four technical replicates, at least three independent experiments were conducted for each treatment.

### Metabolic Labeling with Deuterated Probes

Deuterated metabolic precursors were incorporated into the fly food. For larval experiments, third-instar larvae were transferred to food containing either 20% heavy water (D□O), 50 mM [D□]-glucose, or 50 mM [D□]-acetate for 48 hours before hypoxia and room air treatment.

### Stimulated Raman Scattering (SRS) Microscopy

*Drosophila* larval brains were dissected in Schneider’s insect medium, fixed in 4% PFA (Paraformaldehyde) and mounted for imaging. SRS imaging was performed on a custom-built microscope system. An integrated laser system (picoEmerald, Applied Physics and Electronics, Inc.) produced synchronized pump and Stokes laser beams at 80□MHz repetition rate. The Stokes laser (1032.0□nm, 2□ps pulse width, 0.7□nm beam bandwidth) was intensity modulated by an electro-optic modulator at 10□MHz with >90% modulation depth, and the pump laser (tunable from 700 to 990□nm, 2□ps pulse width, 0.7□nm beam bandwidth) was generated by a built-in optical parametric oscillator. The laser beams were spatially and temporally overlapped using two dichroic mirrors and an integrated delay stage and coupled into an inverted laser scanning microscope (Olympus, FV3000). Both laser beams were focused onto the sample through a 25× water objective (Olympus, XLPLN25XWMP, 1.05□N.A.). The transmitted beams were effectively collected by a high N.A. condenser lens (Olympus, oil immersion, 1.4□N.A.), and the Stokes beam was removed by a bandpass filter (CHROMA, ET890/220□m). The remaining pump beam was detected by a large area (10□×□10 mm^2^) silicon photodiode (THORLABS, FDS1010) reversed biased at 64 DC voltage to increase the saturation threshold and reduce response time. The output current was terminated by a 50□Ω terminator (Mini-Circuits, BTRM-50□+) and filtered with a 9.5–11.5□MHz bandpass filter (Mini-Circuits, BBP-10.7+) to reduce the laser noise. Stimulated Raman loss signal in the pump beam was demodulated at 10□MHz using a lock-in amplifier (Stanford Research Systems, SR844 or Zurich instrument, HF2LI). The in-phase signal was sent to the microscope analog interface box (Olympus, FV30-ANALOG) to generate SRS image. For imaging newly synthesized lipids, the laser frequency difference was tuned to the C-D vibrational resonance at ∼2150 cm□^1^. Images were acquired using custom software, and signal quantification was performed using ImageJ/Fiji. The mean SRS signal intensity in the Eaat1 glial cell bodies (identified by co-expressed GFP) was measured and normalized to the control group.

### Statistical Analysis

All quantitative data are presented as mean ± standard deviation (SD). Statistical comparisons between two groups were performed using a two-tailed Student’s *t*-test.

### Nuclei Isolation

For each condition, approximately 300 brains were collected for snRNA-seq from three independent biological experiments, with each experiment consisting of four technical replicates. Following hypoxia and room air treatment, brains were rapidly dissected in ice-cold Schneider’s Insect Medium supplemented with 10% FBS. Tissues were transferred to 1 mL of ice-cold Nuclei EZ Lysis Buffer (Sigma-Aldrich, NUC101) supplemented with 0.1% Triton X-100, RNase inhibitor (0.5 U/μL), and dithiothreitol (DTT, 1 mM). Tissues were homogenized with a Dounce homogenizer (15 strokes with loose pestle, 15 strokes with tight pestle) on ice. The homogenate was incubated on ice for 5 minutes, then filtered through a 40 μm cell strainer. The filtered suspension was centrifuged at 500 x g for 5 minutes at 4°C. The supernatant was discarded, and the nuclear pellet was gently resuspended in 500 μL of Nuclei Wash and Resuspension Buffer (NWRB; 1x PBS, 1.0% BSA, 0.2 U/μL RNase inhibitor). The nuclei were centrifuged again, resuspended in NWRB, and filtered through a 20 μm strainer. Nuclei concentration and quality were assessed using a Countess II Automated Cell Counter with trypan blue staining.

### Single-Nucleus RNA Sequencing

The isolated nuclei suspension was processed using the Chromium Next GEM Single Cell 3’ Reagent Kits v3.1 (10x Genomics) according to the manufacturer’s protocol. Briefly, nuclei were targeted for a recovery of ∼10,000 per sample. Nuclei were partitioned into Gel Beads-in-emulsion (GEMs) for barcoding and reverse transcription. Following cDNA amplification, sequencing libraries were constructed and assessed for quality using an Agilent Bioanalyzer. Libraries were sequenced on an Illumina NovaSeq 6000 platform to an average depth of 50,000 reads per nucleus.

### Computational Analysis

Raw sequencing data were processed using the 10x Genomics Cell Ranger software (v6.0). Reads were aligned to the *Drosophila melanogaster* reference genome (BDGP6.28/dm6), including pre-mRNA to capture intronic reads essential for snRNA-seq analysis ^34^. The resulting gene-barcode matrices were analyzed using the Seurat R package (v4.1). Nuclei were filtered to remove low-quality profiles and potential doublets, retaining nuclei with >200 and <5000 unique genes detected, and <5% mitochondrial gene content. Data from all samples were integrated using Seurat’s reciprocal PCA-based integration workflow. The integrated data were scaled, and principal component analysis (PCA) was performed. The top 30 principal components were used for graph-based clustering (Louvain algorithm, resolution 0.8) and for visualization using Uniform Manifold Approximation and Projection (UMAP). Cell types were annotated based on the expression of known marker genes from the literature. Differential gene expression (DGE) analysis between conditions within specific cell types was performed using the Wilcoxon Rank Sum test with a log-fold change threshold of 0.25 and a Bonferroni-corrected p-value < 0.05. Gene Set Enrichment Analysis (GSEA) was performed using the fgsea R package and Gene Ontology (GO) biological process gene sets.

## Data availability

The sequencing data obtained in this study has been deposited to the NCBI Gene Expression Omnibus (GEO) (http://www.ncbi.nlm.nih.gov/geo/) under accession number GSE334949 and are publicly available as of the date of publication.

## Contributions

Conceptualization: L.S., G.G.H. and D.Z.; methodology: Y.L., D.Z. and Allen Wang; investigation: Y.L., Q.S., D.Z. and A.W.; formal analysis: Y.L. and C.M.; materials and resources: L.S., G.G.H. and D.Z.; funding acquisition: L.S. and G.G.H.; writing–original draft: L.S. and Y.L.; writing–review and editing, L.S., G.G.H., D.Z. and Allen Wang with input from all authors. The authors have declared no competing interest.

## Acknowledgements

We would like to acknowledge the invaluable contribution of Cody Fine at the UCSD Human Embryonic Stem Cell Core Facility for assistance with cell sorting. This work was supported in part by United States National Institutes of Health (NIH) grants R01AG086548, R01GM149976, U01AI167892, R01HL170107, and R01NS111039, UCSD startup funds, and a Sloan Research Fellowship Award to L.S. Additional support was provided by NIH grant R21NS120023 to G.G.H. This publication includes data generated at the UC San Diego IGM Genomics Center using an Illumina X Plus instrument purchased with funding from an NIH Shared Instrumentation Grant (SIG; S10OD026929).

## Supplementary figures

**Figure S1.**
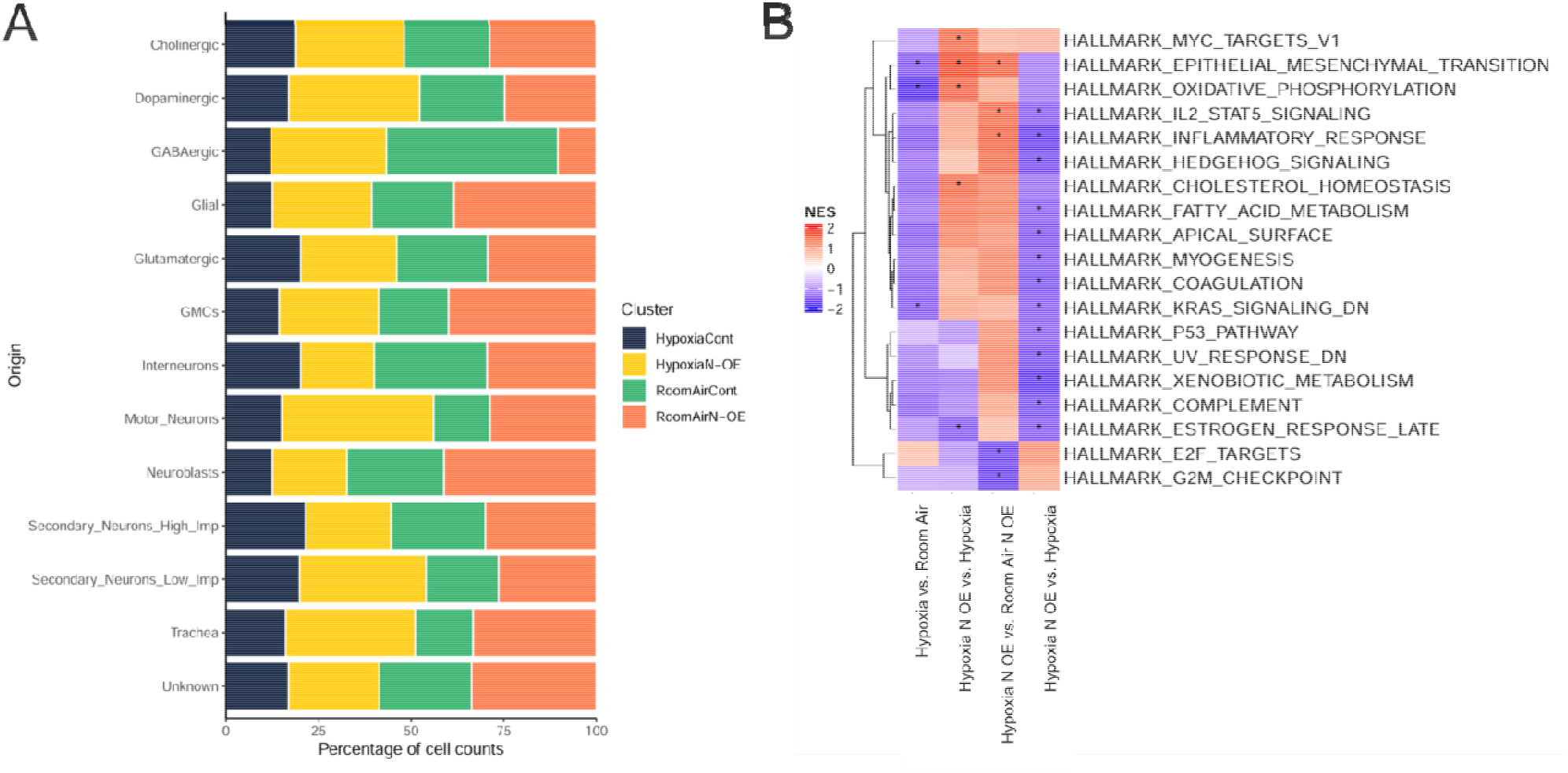
snRNA-seq analysis reveals hypoxia-associated changes in cell composition and transcriptional pathway activity. **(A)** Relative abundance of annotated cell populations across hypoxia, hypoxia N-OE, room-air control, an room-air N-OE clusters. **(B)** Gene set enrichment analysis (GSEA) of hallmark pathways across experimental conditions (room air vs hypoxia; OE vs control). Normalized enrichment scores (NES) are shown for selecte pathways. Hypoxia induces significant changes in pathways related to oxidative phosphorylation, fatty aci metabolism, cholesterol homeostasis, and inflammatory signaling, indicating global metabolic and stress-response remodeling. Red and blue indicate positive and negative enrichment, respectively. Black dots denote significant pathways.

